# Cultural behaviours can be experimentally induced in wild baboons despite constraints on social information transmission

**DOI:** 10.1101/2021.07.12.452028

**Authors:** Alecia J. Carter, Guy Cowlishaw

**Author notes:** Corresponding author (AC).

## Abstract

The formation of culture in animal societies, including humans, relies on the social transmission of information amongst individuals. This spread depends upon the transmission of social information, or social learning, between individuals. However, not all information spreads. To better understand how constraints at the individual-, dyad- and group-level might influence the formation of culture, we experimentally introduced four innovations (novel behaviours) across three troops of wild chacma baboons (*Papio ursinus*). At the individual-level, different phenotypic traits constrained individuals’ use of social information about the innovations. At the dyad-level, we found evidence for social reinforcement and directed social learning affecting who learnt and from whom. Group-level characteristics also limited the diffusion of information, which spread more slowly through social networks that showed less mixing across age classes. Nevertheless, despite these multi-level limitations, the four innovations quickly spread through all the social groups in which they were tested, suggesting that the formation of animal cultures can be surprisingly resilient to constraints on information transmission.

## Introduction

Culture arises through the spread of novel behaviours, or innovations (Castro & Toro, 2004; Kameda & Nakanishi, 2002; McGrew, 1992; Perry & Manson, 2003). This spread depends upon the transmission of social information, or social learning, between individuals (Hoppitt & Laland, 2013). However not all innovations spread (Nishida et al., 2009), and of those that do, the rate of spread can be highly variable (Nishida, 1987). In some cases this may reflect differences in the opportunities for social information transmission (Aplin et al., 2012; Carter et al., 2016). Previous studies have explored how characteristics of individuals, dyads, or groups might influence the spread of innovations (e.g. individuals: (Harcourt et al., 2010; Jones et al., 2017; Thornton & Malapert, 2009); dyads: (Canteloup et al., 2020; Duffy et al., 2009; Schwab et al., 2008); groups (Centola, 2010; Voelkl & Noë, 2010), but none have yet integrated across these three levels of variation to explore how they might relate to one another and how they could combine to affect the emergent patterns of behaviour at the group level. Such an approach is necessary to fully understand the processes that might promote or limit the adoption of innovations and evolution of culture.

Here we assess and integrate the individual-, dyad- and group-level effects on social information transmission about innovations and their consequences on the formation of traditions—socially learnt behaviours adopted by a majority of a group—in a wild primate population. The influences of such effects may be diverse and complex. To begin with individual-level effects, phenotypic constraints may prevent the social learning of an innovated behaviour at three consecutive steps (Aplin et al., 2013; Carter et al., 2016), through limiting either the acquisition of social information, its application, or its final exploitation for benefits (Carter et al., 2016). For example, less social individuals may have fewer opportunities to watch conspecifics and thus acquire social information (Carter et al., 2016), lower-ranking individuals may be unable to access novel resources to apply the information they have acquired (van de Waal et al., 2013), and younger individuals may be too weak to successfully execute an action to exploit the information they have applied (Rapaport & Brown, 2008). Constraints at any one of these three steps may have important implications for the spread of innovations. Thus, for instance, if an individual’s social phenotype prevents the acquisition of information, as seen in spatially peripheral chacma baboons (*Papio ursinus*) (Carter et al., 2016), or the application of social information after its acquisition, as seen in low-status rhesus monkeys (*Macaca mulatta*) avoiding the performance of a socially-learnt task in the presence of high-status monkeys (Drea & Wallen, 1999), this could stall its further transmission to other group members.

In the case of dyad-level effects, the traits of the individual providing the social information (the demonstrator) may influence the willingness of other individuals to learn that information. This is because individuals may preferentially attend to—and acquire information from—particular types of individuals, even if relevant social information is available from others, a process known as directed social learning (Coussi-Korbel & Fragaszy, 1995). For example, captive chimpanzees (*Pan troglodytes*) preferentially copy older, higher ranking demonstrators (Horner et al., 2010); and wild vervet monkeys (*Chlorocebus pygerythrus*) show selective attention towards female demonstrators, resulting in a greater rate of information application in groups with female demonstrators (van de Waal et al., 2010). Such preferences for specific sources of social information could limit the flow of information through a group (Horner et al., 2010) leading to dyadic-level constraints on the spread of innovations (Canteloup et al., 2020).

Finally, at the group level, information transmission may be constrained by structural properties of the social network. Highly clustered networks are predicted to impede information flow as information gets “trapped” in local clusters (Weng et al., 2013), whereas information may spread more rapidly through social networks that are denser—those that have more connections between individuals in the network (Abrahamson & Rosenkopf, 1997). For example, in humans, information was shared more rapidly on the denser Digg social network than the less dense Twitter network (Lerman & Ghosh, 2010). Similarly, regularities within the network may affect information flow. In particular, if individuals with similar phenotypes tend to associate together (positive assortment, or “homophily”), this may preclude others from obtaining social information if they are not associated with the information generators; conversely, “heterophily” may facilitate the transfer of information between information generators and non-generators (Carter et al., 2015). Thus, the diffusion of innovations may be sensitive to network structure at the group level.

The aim of this study is to investigate experimentally how constraints on social information transmission at the individual-, dyad-, and group-level might limit the emergence of cultural traditions. To carry out this investigation, we experimentally created innovations through the presentation of a novel object and three novel foods to three troops of wild baboons, and then assessed the constraints on information flow about these innovations, and their pattern of spread through the troops. In the process, we also test a series of hypotheses about the different mechanisms that might limit the spread of social information at the individual-, dyad-, and group-level. Baboons (*Papio* spp.) are an ideal taxon in which to carry out this study. Like many social species, baboons learn socially (e.g. chacma baboons, *P. ursinus*: (Cambefort, 1981; Carter et al., 2014)), are innovative (chacma baboons: (Marais, 1969); guinea baboons, *P. papio* (Beck, 1973)), and have the capacity to form traditions (e.g., olive baboons, *P. anubis*: (Sapolsky & Share, 2004; Strum, 1975, 1983); guinea baboons: (Claidière et al., 2014)), but baboon cultures are rarely reported.

## Results

### Experimental diffusion of information in baboon groups

To quantify the spread of social information through baboon troops, we introduced four stimuli to small groups of baboons in up to three troops, J, L and M troops, ranging in size from 19 to 67 individuals, while they foraged naturally. One stimulus was a novel object (a brightly coloured cardboard gift box) containing a highly preferred, familiar food, maize kernels. The other three stimuli were novel foods: dried apricot, dried guava and dried mango (Fig. 1). Through these stimuli, we aimed to track the diffusion of the behavioural innovations of taking corn from the gift box, and eating the novel foods, respectively, into the troops. Each stimulus was presented over multiple trials in each troop, in order to provide multiple opportunities for troop members to observe a conspecific interacting with the stimulus. In total, we performed 53 and 46 gift box trials; 150 and 118 apricot trials; 102 and 76 guava trials in J and L troops, respectively, and 61, 50 and 36 mango trials in J, L and M troops, respectively.

**Fig. 1.**
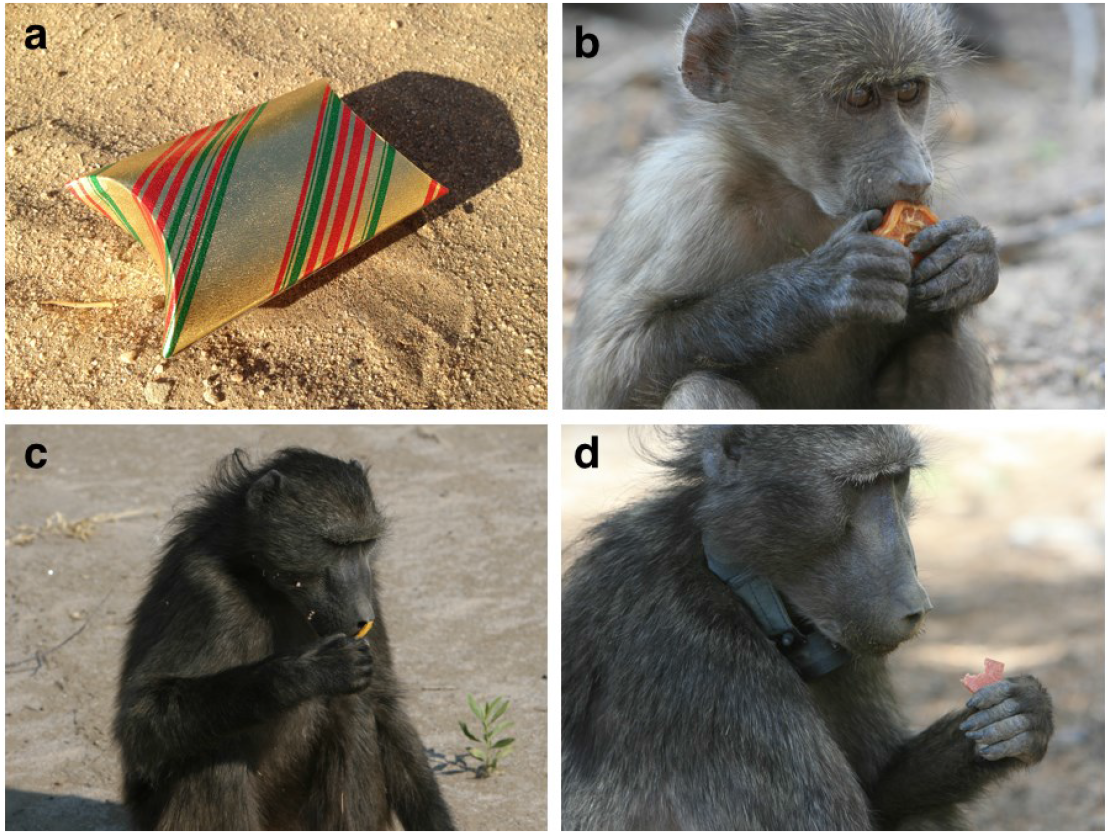
The stimuli used in the diffusion experiments. Shown are the four stimuli: (a) the brightly coloured cardboard gift box that contained approx. 20 corn kernels; (b) an infant baboon consuming the dried apricot; (c) a juvenile male inspecting the dried mango; and (d) an adult female consuming the dried guava.

### Individual-level constraints on information transmission

At the individual level, social information use is a three-step process involving the acquisition of information, its application, and successful exploitation (Carter et al., 2016). To quantify the limitations on information transmission at each of these steps, we compared the characteristics of those individuals who passed through each stage with those that did not across all four diffusions. We focussed on four traits that have previously been linked to social information use across a variety of taxa (including baboons: Carter et al., 2016), namely an individual’s sex (great tits, *Parus major*: Aplin et al., 2015; chimpanzees: Watson et al., 2018), dominance rank (black-capped chickadees, *Poecile atricapillus*: Jones et al., 2017), age (great tits: Aplin et al., 2015; meerkats, *Suricata suricata*: Thornton & Malapert, 2009), and boldness (barnacle geese, *Branta leucopsis*: Kurvers et al., 2010). To estimate the individual characteristics of those who collect social information, while controlling for their opportunities to do so given those with whom they associate, we employed a variant of network based diffusion analysis (NBDA): order of acquisition diffusion analysis (OADA) (Franz & Nunn, 2009; Hoppitt et al., 2010). These analyses are primarily used to determine whether a behaviour is socially learnt by comparing the probability an individual will perform a new behaviour with the social connections an individual has with others who already express the behaviour. We assessed the potential influence of our four traits on whether individuals acquired information by watching a demonstrator, whilst controlling for individuals’ propensities to be near particular others. We found that bolder and lower-ranking individuals were more likely to watch others handling the stimuli (Table 1). In addition, in line with previous findings in this population (Carter et al., 2016), the results of the OADA confirmed that social information was acquired unequivocally through social connections in the proximity network (unbounded *S* = 64.58, ΔAIC_C_ = 293, Likelihood ratio = 295.18, *p* < 0.001).

**Table 1.** Parameter estimates of individual-level variables of the OADA model for asocial effects on information acquisition during the experimental diffusions.

| Individual-level variable | $\beta$ | S.E. | Z | p |
| --- | --- | --- | --- | --- |
| Sex: male<sup>1</sup> | 0.04 | 0.08 | 0.53 | 0.60 |
| Rank | -0.37 | 0.10 | -3.63 | 0.0003 |
| Age class: juvenile<sup>2</sup> | 0.09 | 0.07 | 1.26 | 0.21 |
| Boldness | 0.002 | 0.0008 | 3.13 | 0.002 |

Presented are the individual-level variables, the estimates ( $\beta$ ), the standard errors of the estimate (S.E.), the Z values and p-values.
<sup>1</sup> Reference category: female
<sup>2</sup> Reference category: adult

We next investigated the characteristics of the individuals who applied and exploited social information, using a generalised linear mixed-effects model in each case with the same four individual predictors tested for information acquisition plus an interaction between age and boldness (see Materials and Methods). We found that the probability an individual applied social information, i.e. handled a stimulus after having seen another individual handle it (Table 2), was influenced by the age-boldness interaction effect. Juveniles, regardless of their boldness, were more likely than adults to handle the stimuli after having seen a demonstrator with it, whilst bolder adults were more likely to handle the stimuli after watching a demonstrator with it than shyer adults. In contrast, the probability that an individual successfully exploited the social information they had applied, i.e. obtained corn from the gift box and eat the novel foods, thereby solving the tasks, was influenced by its sex and, possibly, rank, although the latter failed to achieve statistical significance (Table 2). Primarily, male baboons were more likely to solve the tasks once they had handled them after watching a demonstrator.

**Table 2.** Parameter estimates of the reduced models for individual-level traits contributing to differences in information application and exploitation during the experimental diffusions.

| Response | N<sub>obs</sub>, N<sub>ind</sub> | Predictor | B | S.E. | Z | p |
| --- | --- | --- | --- | --- | --- | --- |
| *Information application*<sup>1</sup> | 268, 120 | Intercept | 1.31 | 0.79 | 1.67 | 0.10 |
|  |  | Age class: juvenile<sup>2</sup> | 2.05 | 0.76 | 2.71 | 0.01 |
|  |  | Boldness | 0.39 | 0.19 | 2.06 | 0.04 |
|  |  | Age class:<br>juvenile*boldness | -0.39 | 0.19 | -2.02 | 0.04 |
| *Information exploitation*<sup>1</sup> | 279, 134 | Intercept | 0.82 | 0.80 | 1.03 | 0.30 |
|  |  | Rank | -1.40 | 0.81 | -1.72 | 0.09 |
|  |  | Sex: male<sup>3</sup> | 1.86 | 0.55 | 3.38 | <0.001 |

Presented is the response, the numbers of observations ( $N_{\text{obs}}$ ) and individuals ( $N_{\text{ind}}$ ), the prediction, the estimate ( $\beta$ ), the standard error of the estimate (S.E.), the Z value and p-value. Note that the sample of individuals that exploited the task is larger than the number that handled the stimulus because several individuals' boldness could not be estimated in 2017, and are thus missing from this sample, whilst boldness dropped out of the information exploitation sample during the analysis, increasing the number of individuals.
<sup>1</sup> Coded as 0 for did not handle/solve the stimulus and 1 for handled/solved the stimulus
<sup>2</sup> Reference category: adult
<sup>3</sup> Reference category: female

### Dyad-level constraints on information transmission

To test for directed social learning, we used a variant of network based diffusion analysis (NBDA; Franz & Nunn, 2009; Hoppitt et al., 2010) that allowed us to overcome the disadvantages of more traditional methods used to determine evidence for directed social learning (see Materials and Methods). Our approach, based on dynamic order of acquisition diffusion analysis (OADA), differentially weights observations between particular demonstrators and observers more heavily than others, rather than assuming all demonstrators were equally informative to all observers (Canteloup et al., 2020). We tested eight social transmission hypotheses using OADA: the first three consider social learning strategies which do not involve directed social learning, while the latter five assume directed social learning according to five different demonstrator characteristics. In each case, an animal was considered to have socially learned the task when it obtained corn from the gift box or ate the novel foods after watching a demonstrator. First, we tested whether there was evidence of social learning about the stimuli using the standard, static OADA (Hypothesis 1; H1) compared to asocial learning (H0). The static OADA tested whether the rate at which naïve individuals acquired information from informed individuals was proportional to the social connections they had with informed individuals in the network. We chose a social network based on proximity for the static network (individuals within 10 m), as this has previously been shown to determine information transmission in this population (Carter et al., 2016). Second, we tested two versions of the dynamic OADA that compared whether having witnessed another individual once with the stimulus was enough to subsequently change the observers’ behaviour (dynamic: binary OADA; H2) and another that tested whether individuals were more likely to change their behaviour the more times in total they had seen others handling the stimulus (dynamic: cumulative OADA; H3).

In addition, we tested five hypotheses about directed social learning. These hypotheses involved the traits that were tested at the individual level, and for which there is support for directed social learning in a range of taxa. First, we tested whether adults were more likely to acquire social information provided by other adults (directed: age class; H4) (Barrett et al., 2017; Coelho et al., 2015; Galef & Laland, 2005; Huffman & Hirata, 2003; van de Waal et al., 2010), while juveniles learn equally from both adults and other juveniles. We next tested whether baboons demonstrated a prestige bias, that is, whether they more heavily weighted social information provided by higher-ranking individuals than lower-ranking individuals (directed: rank; H5) (Canteloup et al., 2020; Coelho et al., 2015; Horner et al., 2010; Kendal et al., 2015). Next, we tested whether the baboons were more likely to acquire social information from individuals with whom they had closer social bonds and may thus tolerate in closer proximity than others (directed: tolerance; H6) (van Schaik, 2003). Finally, we tested two hypotheses about the sex of the demonstrator: whether baboons preferentially learnt from the philopatric sex (females in baboons) (van de Waal et al., 2010) or from the same sex (Barrett et al., 2017).

Our OADAs show strong evidence of social learning in the baboons, where all of our alternative hypotheses (H1-H9) gave a better fit to our observations than the null hypothesis of asocial learning (H0). (Table 3; all ΔAIC_C_ > 12.0). Our results found support for both non-directed cumulative learning (H3) and directed learning according to age class (H4), with the former showing a marginally better fit to the data. We found little support for non-directed binary learning (H2) or other forms of directed learning (H5-H8), although all social learning models substantially outperformed the asocial learning null model (H0) (Table 3). This suggests that watching individuals interacting with the stimuli more frequently was enough to predict which individuals would solve the tasks themselves at a later point, particularly if those individuals who were watched were adults.

**Table 3.**
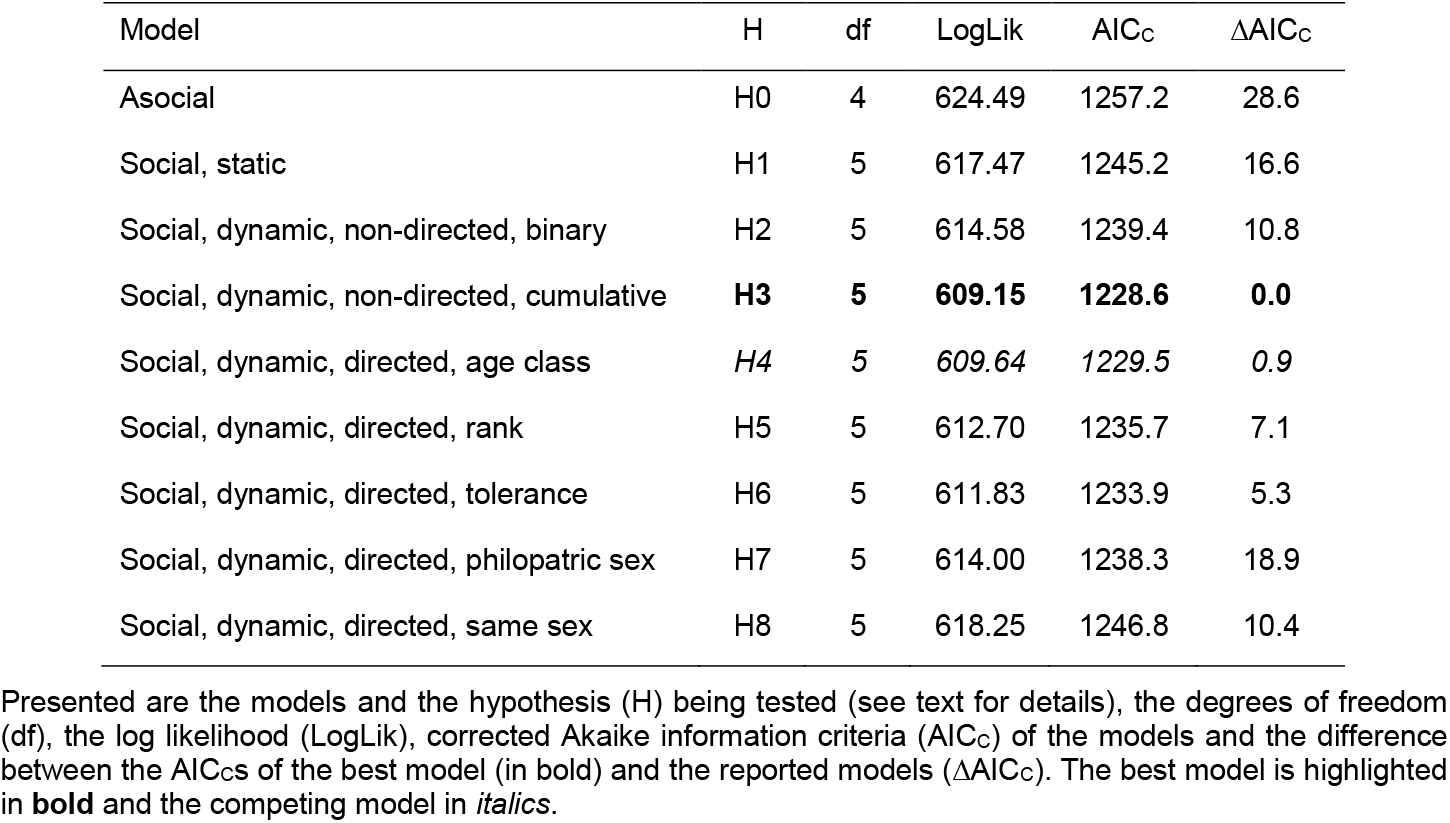
Comparisons of the OADA models with social transmission versus the asocial learning models.

### Group-level constraints on information transmission

To assess whether group characteristics could affect the speed of information diffusion, we calculated a measure of the group “learning rate” for each diffusion, based on the difference, *D*, between the rates with which individuals first had an opportunity to solve the task (first discovered/handled it, whichever came first) and those with which individuals first solved the task (Fig. S1). Because this value is counter-intuitive (larger values = slower diffusions) we calculated 1/*D* as the learning rate. We analysed only the novel-food diffusions as these were directly comparable (all stimuli were included together in the preceding analyses, where it was possible to statistically control for differences between stimuli using random effects). This resulted in seven diffusions across the three troops. Although this number is small, to our knowledge, it represents the most comprehensive number of naturalistic diffusions collected for groups of wild animals.

We tested two general hypotheses about whether the structure of the social networks determined the rate of information flow through them. First, we asked whether two structural properties of the network, specifically transitivity and density, were correlated with the ‘learning rate’. Transitivity, or the clustering coefficient, measures the proportion of triads in a network that are fully connected, that is, how many friends of friends are friends. Networks with higher transitivity have more cliques. Density measures the proportion of all the possible connections among individuals in a network that are actual connections. We predicted that information would transmit more rapidly through less transitive and denser networks, because information would get “trapped” within cliques of highly transitive networks and there would be more opportunities for highly connected individuals to observe others with the stimuli in denser networks, respectively [17,24]. However, there was a strong positive correlation between network transitivity and density in our study troops (Spearman rank correlation test: *S* = 0, ρ = 1, *p* <0.001) so we analysed only network density. We found that the learning rate was unrelated to network density (*S* = 48; ρ = 0.14, *p* = 0.78).

Second, we asked whether the degree of homophily in the networks according to age class, sex and rank was correlated with the learning rates. Homophily is a preference for individuals to be positively assorted in a network i.e. to associate with others of similar phenotypes. We predicted that greater homophily for age class, sex and rank in the networks would result in slower information diffusion. This is because innovative phenotypes, such as juveniles and males (Carter et al., 2014), may preferentially associate with others of similar phenotype and not with those who need to acquire social information for information diffusion to occur (Carter et al., 2015). In line with this prediction, we found that greater homophily for age class was correlated with learning rates such that information diffused more quickly through less assorted networks (Spearman rank correlation test *S* = 106, ρ = −0.89, *p* = 0.01; Fig. 2). Homophily for sex and rank did not correlate with learning rates (Spearman rank correlation tests, sex: *S* = 64; ρ = −0.14, *p* = 0.78; rank: *S* = 72; ρ = −0.29, *p* = 0.56.

**Fig. 2.**
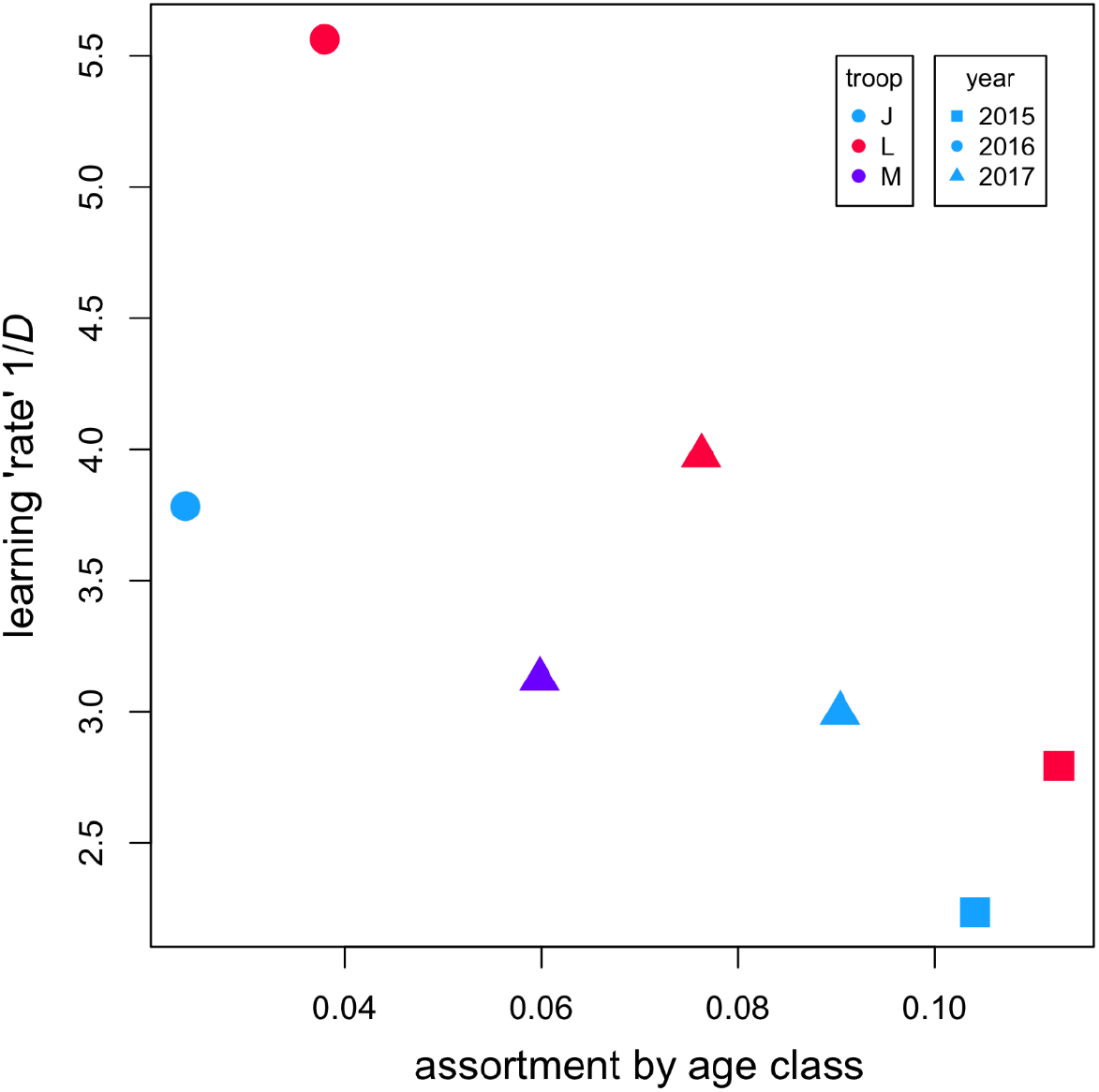
Learning rates for each novel food diffusion were correlated with the homophily in the network. Across the 7 diffusions, the learning rates were correlated with each troop’s assortment by age class in the social network. Each point represents one diffusion, and each troop’s contributions are indicated by different coloured points (see “troop” legend) with each year shown with a different symbol (see “year” legend). Each year equates with the following novel food: 2015: dried apricot; 2016: dried guava; 2017: dried mango.

### Experimentally-induced traditions in wild baboons

Despite the identification of multiple constraints on information transmission, the four innovations quickly spread through each troop. The observed patterns of spread suggest the potential formation of traditions, according to the three criteria used to define traditions (Fragaszy & Perry, 2003). First, a proportion of the group must adopt the new behaviour. In the majority of our diffusions, more than half of the tested individuals in each troop learned to solve the task (median = 57%, range 38-70% of group members) (Fig. 3). These proportions were high given the short timeframe over which the experiments were conducted (2.5-3.5 weeks). Second, the behaviour must be transmitted through social learning. The results from our OADA models indicated that the tasks were socially learnt: all of the social learning models we tested provided a better fit to the data than the asocial learning model (Table 3). Third, the behaviour must persist through time. We were only able to test persistence for one of the novel foods, but in this case persistence was confirmed. In 2016, we re-tested 47 of the 48 individuals who had eaten the apricot in 2015 (one individual died and thus could not be tested) and found 45 of the 47 (96%) persisted in apricot eating. Altogether, the patterns we observed indicated the rapid spread of four socially learned novel behaviours, at least one of which became a tradition. These findings are in line with previous findings of social learning in our population of baboons (Carter et al., 2014) and others (Cambefort, 1981), and mirrors the ‘rapid’ diffusion of information described in other experimental diffusions of information (Aplin et al., 2015).

**Fig. 3.**
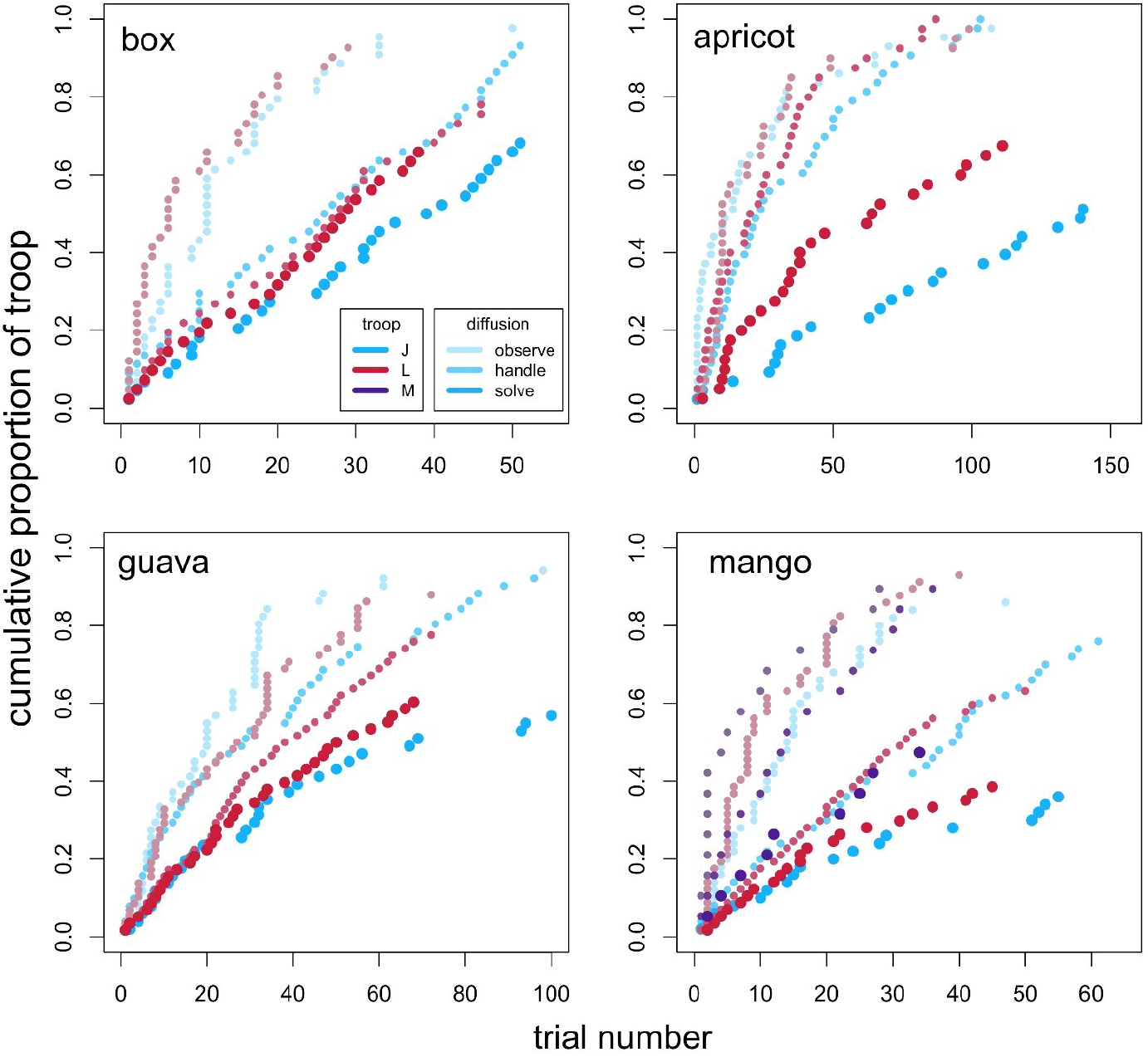
Cumulative proportions of the study troops that had observed others handling the stimuli, had an opportunity to handle the stimuli themselves and solved the tasks. Shown are the cumulative proportions of the study troops, J (blue points), L (red points) and M (purple points, see legend), that observed others handling the stimuli (observe; small points, lowest saturation in colour, see legend for J troop’s change in colour saturation), that handled the stimuli themselves (handle; small, medium saturation) and that solved the tasks (solve; large, highest saturation) for each of the four stimuli, named in each panel, at each given trial. Note that these values reflect the cumulative distributions of individuals’ involvement in the experiments i.e. the trials in which individuals first observed and handled the stimuli and solved the tasks, and do not directly reflect social information acquisition, application and exploitation (see text for details). Note also that the x axes are not on the same scale between stimuli.

## Discussion

The formation of culture depends on the transmission of innovations among members of a group. Constraints on information transmission will thus likely have consequences for the formation of culture. Previous studies have explored such constraints at three different levels: individuals, dyads, and groups. However, these have previously been treated in isolation from one another, and even then rarely explored in wild populations (but see (Aplin et al., 2015; Canteloup et al., 2020) for important exceptions). In this study, we experimentally introduced behavioural innovations in wild groups of chacma baboons, and assessed the influence of individual-, dyadic-, and group-level constraints on the diffusion of these innovations across groups. We found constraints at each level, but these were insufficient to prevent the socially-learned spread of the innovations.

At the individual level, we assessed three steps in the information use sequence: information acquisition, application, and exploitation (following Carter et al., 2016). We found a suite of traits associated with lower use of social information: dominant and shyer animals were less likely to acquire that information, shyer adults were less likely to apply it, and females less likely to exploit it. These patterns reflect a variety of possible mechanisms: dominant animals may have less need to learn about novel foods because they can monopolise food patches and sustain a high-quality diet with familiar foods (Marshall et al., 2012; Post et al., 1980); shyer animals may be less motivated to acquire and, in the case of adults, apply knowledge about novel foods due to their neophobia (Carter et al., 2014); and females may be less motivated to consume novel foods because of their lower food requirements (adult males are nearly twice as large as females) and potential sensitivities to new foods during pregnancy (Mckerracher et al., 2015). These results are generally in line with our previous work demonstrating sequential phenotypic constraints on social information use (Carter et al., 2016), for instance in the finding that females and shyer individuals are less likely to complete the information use sequence. However, there were also two differences, related to age and dominance, that most likely reflect the contrasting nature of the information tested between the two studies: our previous diffusions involved ephemeral information about the location of a high-reward food patch, whereas the present diffusions involved more enduring information about the palatability of potential new foods. We found no age differences in the former, as might be expected given the universal value of that information, but in the present study juveniles were more likely to apply information, presumably reflecting their greater inquisitiveness and interest in interacting with novel foods (Carter et al., 2014; Krueger et al., 2014; Pereira & Fairbanks, 2002). Similarly, we would expect dominant animals to capitalise on ephemeral information about the location of high-reward food patches but subordinates to capitalise on more enduring information about the potential palatability of new foods (e.g. Katzir, 1983). Notably, both dominants and subordinates appeared equally motivated to acquire information about high-reward food patches (the rank difference in that sequence occurred only in exploitation), but subordinates were more likely to acquire information about novel foods (in the present sequence) presumably reflecting their greater potential to benefit from the latter. Such patterns of disinterest by adults and dominant animals, as well as by females and shyer animals, may impede the diffusion of enduring information about potential new foods within a group.

At the dyad level, we found that the cumulative exposure to others handling the stimuli was the best predictor of the individual solving the novel behaviour. Such a result is in line with how wild white-faced capuchins (*Cebus capucinus*) learn to extract seeds from the fruit of *Luehea candida* (Perry, 2009). Likewise, humans are more likely to adopt a novel behaviour if they observe it has been adopted by two or three others in their network rather than by just one (Centola, 2010). This phenomenon is known as *social reinforcement* (Centola & Macy, 2007). To understand this process, it is useful to distinguish between simple and complex contagions. ‘Contagion’ refers to what is being passed between two individuals and could include, for example, information or a virus (Centola & Macy, 2007). Simple contagions require only a brief contact with one knowledgeable (or infected) individual to be passed on and, in humans, could include urban legends or Sars Cov-2. Mathematical models that simulate information diffusion in different theoretical networks have previously highlighted the ‘strength of weak ties’ for predicting rapid diffusion when the contagion is simple (Granovetter, 1977). This is because weak ties can bridge distal parts of a network that may otherwise not be connected. Complex contagions, conversely, require individuals to be exposed to multiple known individuals to adopt a behaviour. That is, changing an individual’s behaviour requires social reinforcement from multiple actors (Centola & Macy, 2007). Experimental studies in humans show that clustered networks—where an individuals’ connections are also connected themselves—facilitate the transmission of complex information because individuals are more likely to be exposed to multiple sources of the information (Centola, 2010). Our finding that the best predictor of dyadic learning was the number of baboons observed others handling the stimuli may provide the first experimental evidence that social reinforcement could play an important role in the diffusion of innovations through animal social networks. Furthermore, it suggests that additions to an established diet are complex contagions. Importantly, social reinforcement is distinct from majority bias and conformity, i.e., a tendency for individuals to change their behaviour to match the behaviour of the majority of the local population (Aplin et al., 2015; Luncz & Boesch, 2014; van de Waal et al., 2013), in three ways: the behaviour being adopted does not require an individual to choose between two (or more) options; the behaviour does not have to be performed by the majority for it to be adopted; and the process describes the diffusion of a behaviour through a population, rather than individuals’ choices that maintain an established cultural behaviour. Social reinforcement presents a new mechanism to better understand when and where innovations will spread in animal societies.

In addition to finding support for social reinforcement, we found that directed social learning by age-class was a competing model for dyadic information transmission. Adults tended to be more likely to solve the task if they had seen another adult performing it. A preference for older demonstrators is relatively common in social learning [sticklebacks, *Pungitius pungitius*: 13; white-faced capuchins: 40; humans: 55], although it may reflect a correlation with proficiency [tufted capuchins, *Sapajus* spp.: 39] or be entirely absent [vervet monkeys: 15; chimpanzees: 41]. Where present, such a preference is likely to impede the diffusion of innovations in cases where juveniles are more likely to be innovators [e.g. foraging techniques in white-faced capuchins: 52]. Such a preference for adult demonstrators may arise through adults’ selective attention towards other adults’ behaviour (van de Waal et al., 2010); selective use of that information after its acquisition (Carter et al., 2016); or a combination of both. Overall, our analysis at the dyad level suggests that constraints on information diffusion may arise when individuals have fewer opportunities to watch others, and when individuals have fewer adults to watch.

At the group-level, we found that diffusions were slower when social networks were assorted by age, i.e., when adults and juveniles associated with others of the same age class. Such a pattern is consistent with our observations of age effects at both individual and dyad levels, where juveniles are more likely to apply information but adults are more influential demonstrators to other adults. If there were no age effects at the individual or dyad level then we would expect no influence of age assortativity at the group level. But the observed age effects suggests that a more heterogeneous distribution of adults and juveniles promoted the diffusion because it allowed some adults to learn by observing juveniles applying information and allowed other adults to subsequently learn from those adults. In a simulation study of information transmission through real primate networks, information was more likely to persist in networks in which there was heterophily for network strength i.e. individuals with more connections preferentially associated with individuals with fewer connections (Voelkl & Noë, 2010). In contrast, an experimental study in human social networks showed that homophily *promoted* the adoption of a novel health behaviour in an online network (Centola, 2011), probably because humans are more likely to be influenced by others who have similar traits to themselves. We might also have obtained a positive effect of homophily if the same age class promoted information transmission at both individual and dyadic levels rather than the opposing age effects that we observed. Network assortativity effects were not found for sex or rank despite each of these traits influencing information transmission at the individual level. This might suggest that mechanisms at the dyad level are more important for learning rates at the group level, or that a combination of both individual- and dyad-level effects are necessary to produce an assortativity effect across the group.

Despite the identification of constraints on social information transmission at the individual-, dyad-, and group-level, we were able to successfully induce the spread of four innovations through social learning in our study groups, at least one of which became a behavioural tradition. The most likely explanation for our success is that the nature of the information involved in our diffusions was favourable to its uptake. Considering the information characteristics that are likely to influence ease of transmission, namely quality, relevance, complexity, congruence, and quantity (Duboscq et al., 2016), our tasks involved stimuli that scored highly in each case: our stimuli involved a reliable food reward from a simple and largely natural stimulus presented on multiple occasions. It may well be that tasks involving less favourable stimuli in any one of these categories would have failed to overcome the constraints. Nunn *et al*. (Nunn et al., 2009) have similarly shown that tasks that are less beneficial are less likely to become adopted in a simulated population. Nevertheless, our findings highlight that the diffusion of new innovations may be sufficiently resilient to overcome a variety of constraints on social information transmission, and suggest that when the characteristics of the information are favourable then the emergence of culture may be inevitable.

## Materials and Methods

### Study area and study species

We performed diffusion experiments over three years (2015-2017). The study subjects were the individually-recognisable members of three habituated troops (J, L and M) of wild chacma baboons at Tsaobis Nature Park, Namibia (15° 45’E, 22° 23’S). Data were collected over 2-3 month periods during the austral winter. Two troops, J and L, were followed over the 3-year study and, following the fission of J troop in 2016, the recently-formed M troop was followed for 2017. Across these seven troop-years, we sampled a median of 50 individuals (range 19-61), representing a median proportion of 0.89 of the entire troop (range 0.65-1), in each diffusion. The median number of females and males was 24 and 27 (ranges 12-29, 10-32) and the median number of adults and juveniles was 22 and 28 (ranges 10-27, 11-40), respectively. Age classes were determined by menarche in females (~4-6 years of age) and canine eruption in males (~7-9 years); individuals were considered as juveniles before these events and adults after. Dominance ranks were assessed through *ad libitum* aggressive interactions recorded during each field season and calculated using the I&SI method (de Vries, 1998) implemented in Matman 1.1.4 (Noldus Information Technology, 2003). All displacements, supplants, threats, chases and attacks for which we could identify both actor and recipient were recorded. If more than one dominance behaviour occurred in one event, such as a threat followed by a chase, only one interaction was recorded. The dominance hierarchies were strongly linear (Landau’s corrected linearity index *h*’, *p* < 0.001 in all cases). Dominance rank was expressed relatively to control for group size using the formula 1-[(1-*r*)/(1-*n*)] where *r* is the individual’s absolute rank and *n* is the group size, and ranges from 0 (lowest rank) to 1 (highest rank). Further details on the study periods, sample sizes, and dominance data are provided in Table S1. Personality was indexed by boldness, estimated by presenting individuals with a novel food while foraging naturally alone and quantifying the time that individuals spent investigating—handling and smelling—the novel foods up to 120 s. The investigating time was used as an individual’s boldness score as it has convergent validity with observers’ ranks of individuals’ boldness and is repeatable over three years (Carter et al., 2012a, 2012b). The novel foods were an eighth of an apple dyed blue in 2015, three popped corn kernels in 2016, and a peanut in its shell dyed blue or red in 2017. Individuals who investigated the novel food for longer were considered bolder. We tested individuals’ boldness annually during the study; however, due to the difficulty of testing juveniles when alone, in 2016 and 2017 only 94 of 114 and 70 of 131 total individuals were tested.

### Experimental design

To experimentally diffuse four innovations we introduced stimuli to small groups of baboons while they foraged naturally. In 2015 the stimuli were (i) a red-, green- and gold-striped patterned gift box (pillow box dimensions: 6.6 cm x 6.6. cm x 2.5 cm, shopping-bags4u.co.uk, see Video 1, Fig. 1) that contained approx. 20 kernels of popping corn (maize), a familiar, highly-preferred food; and (ii) commercially-processed dried apricots, a novel, unfamiliar food; in 2016, (iii) a quarter of a commercially-processed dried guava slice, a second novel food; and in 2017, (iv) a 4-5 cm long, 1-2 cm wide strip of dried mango, a third novel food. Very few wild baboons in our population readily eat novel foods when first encountered. We have been using responses to novel foods to determine systematically individuals’ boldness since 2009 (for example, see Carter et al., 2012a, 2012b). Over this 3-year study, the proportions of individuals willing to eat a novel food on first encounter ranged from 0% of the troops (0 of 94 individuals ate cooked corn kernels in 2016) to 29% (20 of 70 individuals ate peanuts from shells dyed red or blue in 2017); on average, 17% of individuals ate the novel foods presented. Furthermore, without having seen others handling the novel foods, individuals spend less time investigating them on subsequent presentations (Carter et al., 2014), suggesting that sequential gathering of personal information is not sufficient for individuals to adopt a novel food into their diet. Novel foods thus present appropriate stimuli with which to assess the diffusion of information through social learning in this species. The troops received the gift box from 27 June-20 July 2015, the apricot from 4-26 September 2015, the guava from 30 July-16 August 2016, and the mango from 18 July-23 August 2017.

One observer (AJC) presented the stimuli within 15 m of subgroups of foraging baboons (that could comprise 2-~10 individuals of the troop) either by placing the stimulus behind her bag whilst pretending to get something from it (gift box) or covertly dropping the stimulus behind her as she was standing/walking in front of or next to a subgroup (novel foods). Two considerations were key: first, that the presentations were representative of naturalistic diffusions of information, such as the adoption of a novel (e.g. alien) species of plant into the diet, and second, that every baboon had at least one opportunity to collect social and/or personal information about the tasks. Whilst we were interested in limitations on social information transmission, at least one individual in each subgroup had to first collect personal information about each stimulus before it could be socially acquired by others. As such, our experiments represented open troop diffusions with no seeded demonstrators (Whiten & Mesoudi, 2008). In addition, subgroups were targeted in such a way as to maximise the numbers of individuals that received opportunities to interact with the tasks, but to avoid frequently presenting the stimuli to the same individuals lest they associate the observers with food, which may have changed their behaviour. As such, during the initial stages of the diffusions, multiple individuals were first given opportunities to collect personal information of the stimuli. This is in contrast to diffusions in captive situations, where there are usually one or two trained demonstrators (e.g. Horner et al., 2010; Whiten & Mesoudi, 2008). Furthermore, whilst all individuals were given opportunities to acquire both personal and social information, not all did so because there was no compulsion either to interact with the stimuli or to look at a demonstrator with the stimulus, respectively. All trials were video recorded to facilitate later data extraction. We recorded the identities of all individuals who (i) looked at an individual handling the stimuli, (ii) handled the stimuli themselves and (iii) solved the task by extracting food from the box or eating the novel food (see Videos 1 and 2). Across the four diffusions, most individuals observed others with and handled the stimuli; in all cases >85% of individuals in a given troop observed and >57% handled the stimuli (Fig. 3, Table S2). Note that the values in Fig. 3 and Table S2 reflect the cumulative totals of individuals’ involvement in the experiments and do not directly reflect social information acquisition, application and exploitation. This is because the cumulative distributions include some individuals who collected personal information before social information [as is the case in all natural diffusions, some individuals will encounter the stimulus before observing others with it], whereas our analyses of individual-level social information application and exploitation were limited solely to those individuals who had acquired social information before interacting with the stimulus.

### Quantifying social connections

We created social networks based on the number of times individuals were observed in close proximity to others. These data were used to build baseline networks for use in the individual- and dyad-level NBDA analyses and to calculate group-level network statistics for the group-level analyses. The proximity data were collected throughout each field season but, to ensure independence from the experimental data, were not collected at the same time as the experimental presentations. We recorded subgroups within each troop between dawn and dusk during all behavioural states. To ensure subgroups were sampled randomly and individuals were sampled evenly (and there was no bias against less sociable individuals), we quantified the subgroups associated with given ‘focal’ individuals chosen randomly from the troop membership. Because the troops can spread over 1 km^2^ and finding particular individuals can be time-consuming, the observer searched for one of the first five individuals on a randomised list to optimise the number of independent subgroups sampled each day. Once a focal individual was found and its subgroup membership quantified, that individual was removed from the list until all remaining individuals had been found and a new list started. If a focal individual had already been recorded in an already-sampled subgroup (i.e., group membership had not changed), that individual was not sampled for an hour to ensure independent data. We are confident each sampled subgroup was independent and as such have not pooled data within arbitrary time periods [*cf*. 66]. Subgroup membership was recorded using an instantaneous scan of the identities of all individuals within a 10 m radius of the focal as we have previously found this social network to best predict information transmission among individuals (Carter et al., 2016). Those without a neighbour within 10 m were recorded as alone in a subgroup size of 1. We collected on average (median) 1089 10 m scans per troop-year (range J: 578-1381; L: 679-1271; M: 416). These were used to build proximity networks at the troop-year level using a simple ratio index (Whitehead, 2008).

### Statistical analyses

#### Individual-level constraints on the social information use sequence

The first stage of the information use sequence is the acquisition of social information, i.e., watching another handling the stimulus. To estimate the individual characteristics of those who collect social information, while controlling for their opportunities to do so given those with whom they associate, we employed network based diffusion analysis (NBDA). NBDA and its variants (Franz & Nunn, 2009; Hoppitt et al., 2010) are primarily used to determine whether a behaviour is socially learnt by comparing the probability an individual will perform a new behaviour with the social connections that individual has with others who already express the behaviour. One advantage of NBDA is that it can incorporate individual-level variables to control for possible sources of individual variation in asocial learning ability. We took advantage of this to estimate not the order that the task was solved, but the order in which individuals collected social information about the stimuli. By doing so, we could estimate the characteristics of those who were more likely to watch others with the stimulus, i.e. collect social information, whilst controlling for the probability that they would be in close proximity to particular others, i.e. controlling for the availability of social information. It is important to understand that this interpretation is not the traditional use of NBDA: for this analysis we were not interested in whether the social network predicted the order that individuals collected social information while controlling for individuals’ characteristics, but which individual characteristics determined social information acquisition while controlling for opportunities to collect social information (i.e., the proximity network). For each presentation, we identified all individuals who watched another handling the stimulus (the demonstrator) and in which order. This resulted in a “diffusion” of social information for each presentation. This is different to the diffusion of the task itself, which occurred over the course of all the presentations as more and more individuals solved the task. We then employed a variant of NBDA, order of acquisition diffusion analysis (OADA), which uses the order in which individuals acquired the information rather than the time that they acquired it (Hoppitt et al., 2010), to estimate the effects of individual-level variables on asocial “watching” ability. We included age class, sex, rank and boldness as individual-level variables. To maximise power, all diffusions were included in the analysis, rather than analysing them separately, and each of the four stimuli was given a unique ID to control for differences between the stimuli. We removed all presentations in which there was no diffusion of information (i.e., no observers of the demonstrator), along with two guava and one mango presentation because an unidentifiable individual for whom we did not have social network data was the demonstrator. This resulted in, for J and L troops respectively, 100 and 86 diffusion events for the apricot, 44 and 39 events for the gift box, and 65 and 52 events for the guava, and, in J, L and M troops, 39, 38 and 29 events for the mango, resulting in a total of 492 diffusions of social information across the four experiments. We used a multiplicative OADA with the respective social networks for the given years of study.

The second and third stages of the information use sequence are the application and exploitation of social information, i.e., the handling of stimuli and the solution of the tasks (getting corn from the box or eating the novel foods), respectively. We compared the characteristics of those individuals that handled and solved the tasks with those that did not using generalised linear mixed models (GLMMs) with a binomial response (handle/solve or not) and the experiment and baboon identity nested within troop as random effects to control for both repeated observations of baboons across the four diffusions and the potential differences in the ease of the task to solve. We included as fixed effects individual age class, sex, rank and boldness. We also included an interaction between age class and boldness, as we have previously found these characteristics to be important for information use in this species (Carter et al., 2014, 2016) and our data suggest that boldness differentially changes with age depending on sex (A. Carter, unpublished data). We used backwards elimination of non-significant terms until we arrived at a minimal model. For information application, we restricted the analysis to individuals who had collected social information and had not solved the task before watching others interacting with it (i.e. the social information collected was still relevant at the time it was collected). This sample included, over the four stimuli, 54 individuals who did not interact with the stimuli—by handling and/or solving it—after watching others with it and 279 who did. This sample will include some individuals who likely would have interacted with the task without having collected social information, but this is representative of natural diffusions. For information exploitation, we restricted our analysis to individuals from the above sample who did interact with the stimuli, i.e. applied social information. This sample included 105 individuals who did not solve the task after watching a demonstrator and 174 who did.

#### Directed social learning

Identifying directed social learning typically requires prolonged experiments where multiple demonstrators are trained to solve different tasks, and the subsequent adoption of each task in the group is examined to determine whether there is a bias towards one, indicating the preferred (type of) demonstrator (e.g. Horner et al., 2010). Although this methodology is rigorous for identifying biases, few demonstrators are used; it usually requires a lengthy training period, followed by a lengthy demonstration period and short test period; and many studies are completed in captivity, where particular demonstrators can be trained away from other group members (but see van de Waal et al., 2010) and the chosen demonstrators may not necessarily reflect the types of individuals who are likely to generate novel information in the wild (Biro, 2011; Carter et al., 2015; van Schaik, 2003).

To test for directed social learning in a way that overcomes these limitations as far as possible, we developed a statistical method to identify constraints on social information use that may arise through directed social learning during naturalistic diffusions by building on a variant of NBDA (Franz & Nunn, 2009; Hoppitt et al., 2010), the dynamic order of acquisition diffusion analysis (dynamic OADA; Hobaiter et al., 2014). Hobaiter *et al*. (2014) extended the static version of OADA, which infers opportunities for information transmission based on rates of association, to a dynamic OADA, which uses a network of actual observations of the transmitted behaviour which is updated at each time step as it diffuses through the group. Dynamic OADA takes advantage of detailed information about who observes whom handling a stimulus, and models its diffusion according to a cumulative total of times individuals have been observed observing others. In this way, it considers temporal dynamics in individuals’ exposure to social information.

We adapted the dynamic OADA to test amongst competing hypotheses about who learns from whom. In a similar manner to Canteloup *et al*. (Canteloup et al., 2020), we differentially weighted observations between particular demonstrators and observers more heavily than others, rather than assuming all demonstrators were equally informative to all observers. In the first case, we calculated an OADA model using the traditional ‘static’ 10 m proximity network (H1: static). Next, following Hobaiter *et al*. (Hobaiter et al., 2014), we then tested two versions of the dynamic OADA. The first compared whether having witnessed another individual once with the stimulus was enough to subsequently change the observers’ behaviour (H2 dynamic: binary OADA). The second tested whether individuals were more likely to change their behaviour the more times in total they had seen any others handling the stimulus, regardless of who they observed (H3 dynamic: cumulative OADA). We also tested five hypotheses about directed social learning. First, we tested whether adults were more likely to acquire social information provided by other adults (H4 directed: age class) (reviewed in Galef & Laland, 2005; Huffman & Hirata, 2003; van de Waal et al., 2010) by weighting observations of juveniles by adults as 0 (i.e. providing no social information), and all other observations as 1 (i.e. providing social information). This is a conservative estimation which suggests that adults do not use any information received from juveniles. Second, we tested whether baboons demonstrated a prestige bias, that is, whether they were more attentive towards social information provided by higher-ranking individuals (H5 directed: rank) (Horner et al., 2010). We did this by weighting the (cumulative) observations by the demonstrators’ ranks. This meant that individuals who watched an individual of rank 0.2 three times would have acquired the ‘equivalent’ social information as an individual who observed an individual ranked 0.6 one time. Again, this is a conservative estimation of directed social learning through a prestige bias. Next, we tested whether the baboons were more likely to learn from individuals with whom they had closer social bonds. We did this by weighting the observation of a demonstrator by the strength of its social connection with the observer, based on the frequencies of close proximity within 5 m to reflect the kind of social tolerance predicted to increase opportunities for social learning (H6 directed: tolerance) (van Schaik, 2003). Finally, we tested two hypotheses about the sex of the demonstrator. In the first case, we tested whether baboons were more likely to learn from the philopatric sex (van de Waal et al., 2010) in the same manner as the age-class model, by weighting observations of females as 1 and observations of males as 0 (H7 directed: philopatric sex). In the second case, we tested whether individuals were more likely to learn from an individual of the same sex (Barrett et al., 2017) by weighting observations of a same-sex demonstrator as 1 and an opposite-sex demonstrator as 0 (H8 directed: same sex). To maximise statistical power, for each of the eight dynamic OADA models the four stimuli were analysed together, with a unique ID assigned to each stimulus to control for differences between them. We determined the best model as the one with the lowest Akaike Information Criterion corrected for small sample size (AIC_C_).

#### Group-level constraints on information diffusion

We calculated a measure of the group “learning rate” for each diffusion by calculating the difference between the rate that individuals first had an opportunity to solve the task and the rate at which they actually solved it. The learning rate was estimated in two steps. First, we calculated the areas under the curves made when the cumulative proportions of the troop (a) discover or handle the stimulus, and (b) solve the task, were plotted against trial number (Fig. S1). To make the different diffusions comparable between troops, the trial numbers were standardised to terminate at one by dividing the trial number by the total number of trials. Second, we calculated the difference in these areas (a – b), interpreting larger values as indicative of slower information diffusion. It is possible that, over the 3-year course of the study, the baboons learnt a generalizable “rule” about dried novel fruits—that they are edible—which could affect the learning rates. That is, the baboons may have needed less social information about each subsequent novel fruit to eat it, and information diffusion became correspondingly faster as the study progressed. However, there was no relationship between learning rate and study year (Spearman rank correlation *S* = 82.46, *p* = 0.28), suggesting that the baboons treated each novel fruit as an equally novel stimulus.

We tested whether the density and assortment of the social networks could determine the rate of information flow through them. Density is a social network metric that calculates the proportion of potential edges in a network that are actual edges: networks have higher density when more of the potential connections between individuals are actual connections. Assortment is the tendency for connected individuals to be similar to each other and ranges from −1 (individuals associate only with others of different traits) to 1 (individuals associate only with others of similar traits), where 0 is indicative of random association according to the trait of interest. We restricted our analyses to the novel food diffusions as these were replicated across the three years of the study and are comparable amongst each other. For each troop in each year, we calculated the network density using the R package sna (Butts, 2008) and assortment by age class, sex and rank using the R package asnipe (Farine, 2013), resulting in a total of 4 network metrics per troop. We did not calculate assortment by boldness as we had missing values in 2016 and 2017, which asnipe cannot handle. To test whether these network-level characteristics predicted the speed of information transmission through the baboon groups, we ran Spearman rank correlations because of the small sample size of diffusions.

### Ethical approval

This study was performed in accordance with the recommendations of the Association for the Study of Animal Behaviour’s Guidelines for the Treatment of Animals in Behavioural Research and Teaching. Our protocols were assessed and approved by the Ethics Committee of the Zoological Society of London (Permit: 041-BPE-0524), and approved by the Ministry of Environment and Tourism in Namibia (Research/Collecting permits 2009/2015, 2147/2016, 2303/2017).

## Acknowledgements

We give a huge thank-you to Carolyn Dugon, Dylan Gomes, Debbie Walsh, and Katie Stone for collecting much of the proximity social network data. We also thank Claudia Martina Luna, who allowed us to complete our field work alongside hers in both 2015 & 2016 and Alice Baniel, who co-directed the 2017 field season with a pregnancy-diminished AJC. We thank the members of the Tsaobis Baboon Project 2015, 2016 and 2017 for collecting many of the interaction data and Alex Lee for help with the dominance hierarchies. We thank Will Hoppitt for explaining how to modify OADA to make it dynamic. AJC was supported in 2015 and 2016 by a Junior Research Fellowship from Churchill College, University of Cambridge. The fieldwork was supported by a Gibbs Travelling Fellowship from Newnham College, University of Cambridge (2015) and a Leakey Foundation Research Grant from Dennis Fenwick and Martha Lewis (2016-2017). This research was carried out with the permission of the Ministry of Environment and Tourism and the Ministry of Land Reform, and in affiliation with the Gobabeb Namib Research Institute. We thank the Tsaobis beneficiaries for permission to work at Tsaobis, and Johan Venter and the Snyman and Wittreich families for permission to work on neighbouring land. This paper is a publication of the ZSL Institute of Zoology’s *Tsaobis Baboon Project*.

## Supplementary Materials for

**Table S1.**
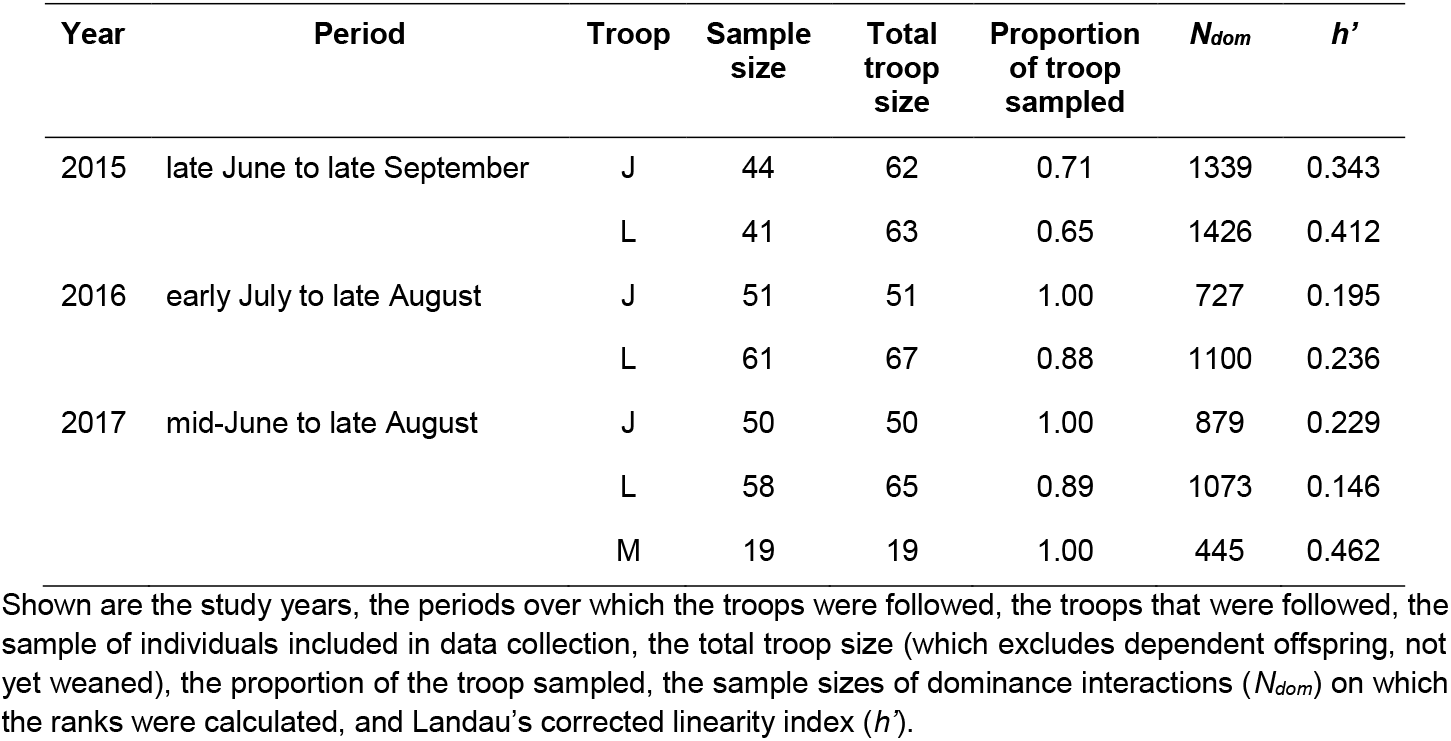
The samples of individuals and data collected over the three year study period.

**Table S2.** The number of times individuals watched a demonstrator, handled a stimulus and solved a task.

| Task | J troop |  |  |  | L troop |  |  |  | M troop |  |  |  |
| --- | --- | --- | --- | --- | --- | --- | --- | --- | --- | --- | --- | --- |
|  | Total cases | Individuals (% total of tested troop) | Range | Median | Total cases | Individuals (% total of tested troop) | Range | Median | Total cases | Individuals (% total of tested troop) | Range | Median |
| *Box* |  |  |  |  |  |  |  |  |  |  |  |  |
| Observed a demonstrator | 128 | 41 (93) | 0-9 | 1 | 162 | 38 (93) | 0-8 | 1 |  |  |  |  |
| Handled the stimulus | 54 | 33 (75) | 0-5 | 1 | 47 | 31 (76) | 0-3 | 1 |  |  |  |  |
| Solved the task | 44 | 31 (70) | 0-5 | 1 | 39 | 28 (68) | 0-3 | 1 |  |  |  |  |
| *Apricot* |  |  |  |  |  |  |  |  |  |  |  |  |
| Observed a demonstrator | 237 | 42 (95) | 0-11 | 5 | 220 | 39 (95) | 0-16 | 5 |  |  |  |  |
| Handled the stimulus | 162 | 36 (82) | 0-14 | 4 | 142 | 39 (95) | 0-9 | 3 |  |  |  |  |
| Solved the task | 82 | 22 (50) | 0-9 | 3 | 82 | 26 (63) | 0-9 | 3 |  |  |  |  |
| *Guava* |  |  |  |  |  |  |  |  |  |  |  |  |
| Observed a demonstrator | 161 | 48 (94) | 0-10 | 3 | 111 | 54 (89) | 0-7 | 2 |  |  |  |  |
| Handled the stimulus | 108 | 45 (88) | 0-5 | 2 | 90 | 53 (87) | 0-3 | 1 |  |  |  |  |
| Solved the task | 60 | 29 (57) | 0-4 | 1 | 59 | 36 (59) | 0-3 | 1 |  |  |  |  |
| *Mango* |  |  |  |  |  |  |  |  |  |  |  |  |
| Observed a demonstrator | 125 | 43 (86) | 1-8 | 2 | 130 | 53 (91) | 1-9 | 2 | 61 | 18 (95) | 1-6 | 3 |
| Handled the stimulus | 77 | 41 (82) | 1-4 | 2 | 70 | 45 (58) | 1-3 | 1 | 42 | 17 (89) | 1-6 | 2 |
| Solved the task | 33 | 19 (38) | 1-4 | 2 | 33 | 22 (38) | 1-3 | 1 | 23 | 9 (47) | 1-8 | 2 |

Presented are the total number of cases that individuals watched another individual handling the stimulus (watched a demonstrator); handled the stimulus themselves; and solved the task (ate the corn kernels or novel foods). Note that these numbers reflect the overall distributions of individuals' involvement in the experiments and do not directly reflect social information acquisition, application and exploitation as individuals (see text for details).

**Fig. S1.**
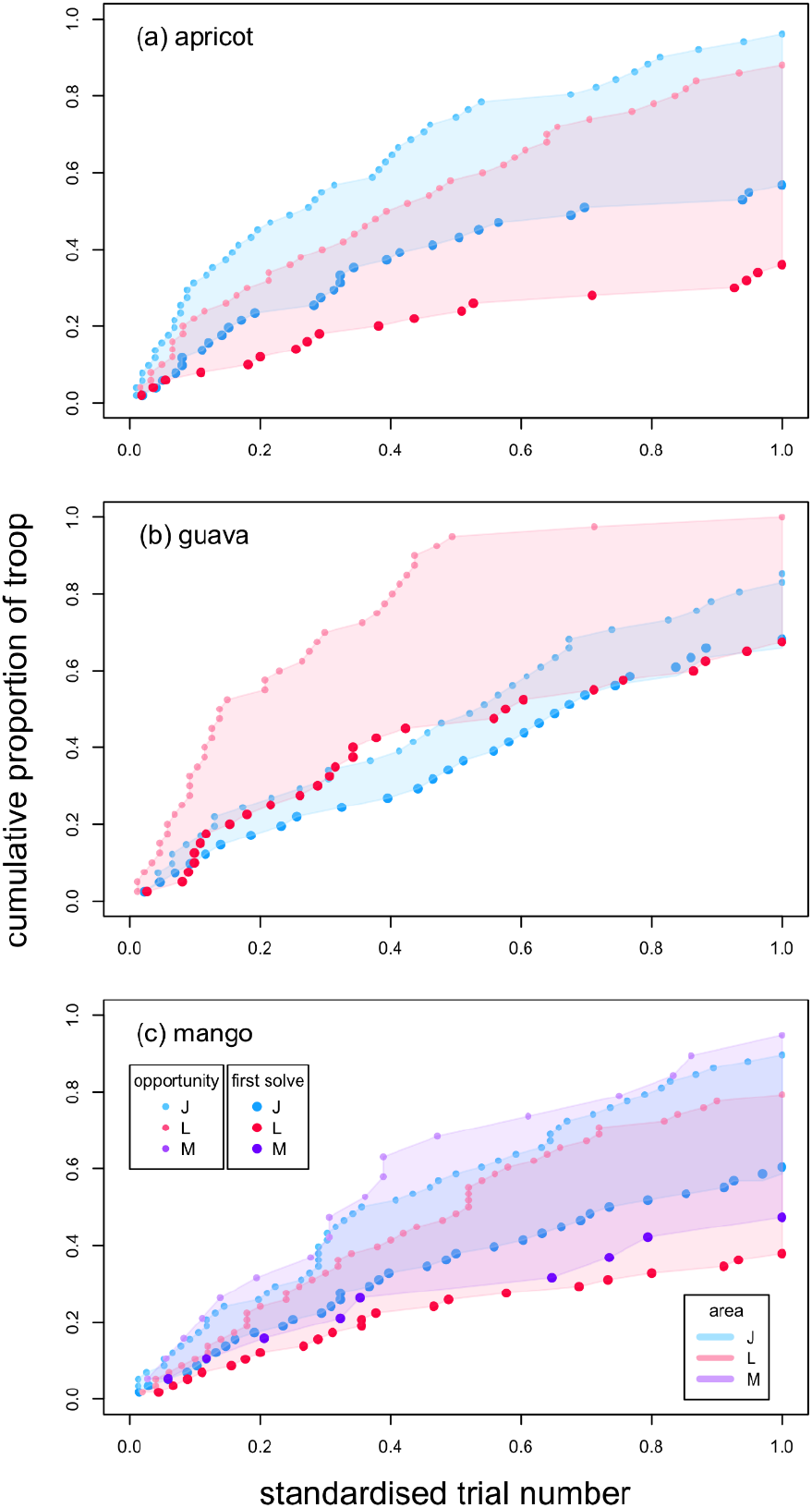
Learning “rates” for each troop for each novel fruit diffusion across three years. Troop-level learning rates for each diffusion, were calculated as the difference, *D* (shaded areas, see “area” legend), between the area under the “solve” curve (larger, darker points, “first solve” legend) and the area under the “opportunity to solve” curve (smaller, lighter points, “opportunity” legend), which was the first time an individual discovered the stimulus or handled the stimulus after watching a demonstrator. The diffusions for each stimulus (a: apricot, b: guava and c: mango) and the shaded areas (*D*) for each troop (see legends) are shown.

## Video S1

### Gift box experiment

The gift box was placed on the ground outside of a large patch of *Salvadora persica* in which several baboons were feeding. Two naïve adult males come near to, but do not see, the gift box. Both a naïve juvenile male and knowledgeable adult female see the box. Both approach when the adult males leave the vicinity; the adult female arrives first and handles the box, observed by the juvenile male. A knowledgeable juvenile female also approaches and observes the adult female with the box. The adult female solves the task, observed by both the juvenile male and female. Three infant baboons approach, and watch the female with the gift box. Both the juvenile male and female interact with the remains of the box after the adult female leaves. Note that, while the video is trained on the individual interacting with the box, the observer is scanning the surrounding area for other baboons.

## Video S2

### Novel food experiment

The novel food, a dried apricot, was placed on the ground between patches of *Prosopis glandulosa*. A naïve adult female approaches and handles the novel food. She solves the task, consuming the apricot. A naïve female approaches and observes the solver. The solver moves away. The naïve adult female approaches the solver again, and is approached and observed by a naïve juvenile male. The juvenile male finds a small piece of apricot dropped by the solver and consumes it. Note that, while the video is trained on the individual interacting with the box, the observer is scanning the surrounding area for other baboons.

## References

Abrahamson, E., & Rosenkopf, L. (1997). Social network effects on the extent of innovation diffusion: A computer simulation. Organization Science, 8(3), 289–309.

Aplin, L. M., Farine, D. R., Morand-Ferron, J., Cockburn, A., Thornton, A., & Sheldon, B. C. (2015). Experimentally induced innovations lead to persistent culture via conformity in wild birds. Nature, 518(7540), 538–541.

Aplin, L. M., Farine, D. R., Morand-Ferron, J., & Sheldon, B. C. (2012). Social networks predict patch discovery in a wild population of songbirds. Proceedings of the Royal Society B: Biological Sciences. https://doi.org/10.1098/rspb.2012.1591

Aplin, L. M., Sheldon, B. C., & Morand-Ferron, J. (2013). Milk bottles revisited: Social learning and individual variation in the blue tit, Cyanistes caeruleus. Animal Behaviour, 85(6), 1225–1232. https://doi.org/10.1016/j.anbehav.2013.03.009

Barrett, B. J., McElreath, R. L., & Perry, S. E. (2017). Pay-off-biased social learning underlies the diffusion of novel extractive foraging traditions in a wild primate. Proceedings of the Royal Society of London B: Biological Sciences, 284(1856). https://doi.org/10.1098/rspb.2017.0358

Beck, B. B. (1973). Observation learning of tool use by captive Guinea baboons (Papio papio). American Journal of Physical Anthropology, 38(2), 579–582.

Biro, D. (2011). Clues to Culture? The Coula- and Panda-Nut Experiments. In T. Matsuzawa, T. Humle, & Y. Sugiyama (Eds.), The Chimpanzees of Bossou and Nimba (pp. 165–173). Springer Japan. https://doi.org/10.1007/978-4-431-53921-6_18

Butts, C. T. (2008). Social network analysis with sna. Journal of Statistical Software, 24(6), 1–51.

Cambefort, J. P. (1981). A Comparative Study of Culturally Transmitted Patterns of Feeding Habits in the Chacma Baboon Papio ursinus and the Vervet Monkey Cercopithecus aethiops. Folia Primatologica, 36(3–4), 243–263.

Canteloup, C., Hoppitt, W., & van de Waal, E. (2020). Wild primates copy higher-ranked individuals in a social transmission experiment. Nature Communications, 11(1), 1–10.

Carter, A. J., Lee, A. E. G., Marshall, H. H., Ticó, M. T., & Cowlishaw, G. (2015). Phenotypic assortment in wild primate networks: Implications for the dissemination of information. Royal Society Open Science, 2(5). https://doi.org/10.1098/rsos.140444

Carter, A. J., Macdonald, S. L., Thomson, V. A., & Goldizen, A. W. (2009). Structured association patterns and their energetic benefits in female eastern grey kangaroos, Macropus giganteus. Animal Behaviour, 77(4), 839–846.

Carter, A. J., Marshall, H. H., Heinsohn, R., & Cowlishaw, G. (2012a). Evaluating animal personalities: Do observer assessments and experimental tests measure the same thing? Behavioral Ecology and Sociobiology, 66(1), 153–160. https://doi.org/10.1007/s00265-011-1263-6

Carter, A. J., Marshall, H. H., Heinsohn, R., & Cowlishaw, G. (2012b). How not to measure boldness: Novel object and antipredator responses are not the same in wild baboons. Animal Behaviour, 84(3), 603–609. https://doi.org/10.1016/j.anbehav.2012.06.015

Carter, A. J., Marshall, H. H., Heinsohn, R., & Cowlishaw, G. (2014). Personality predicts the propensity for social learning in a wild primate. PeerJ, 2, e283. https://doi.org/10.7717/peerj.283

Carter, A. J., Torrents Ticó, M., & Cowlishaw, G. (2016). Sequential phenotypic constraints on social information use in wild baboons. ELife, 5, e13125. https://doi.org/10.7554/eLife.13125

Castro, L., & Toro, M. A. (2004). The evolution of culture: From primate social learning to human culture. Proceedings of the National Academy of Sciences of the United States of America, 101(27), 10235–10240.

Centola, D. (2010). The spread of behavior in an online social network experiment. Science, 329(5996), 1194–1197.

Centola, D. (2011). An experimental study of homophily in the adoption of health behavior. Science, 334(6060), 1269–1272.

Centola, D., & Macy, M. (2007). Complex contagions and the weakness of long ties. American Journal of Sociology, 113(3), 702–734.

Claidière, N., Smith, K., Kirby, S., & Fagot, J. (2014). Cultural evolution of systematically structured behaviour in a non-human primate. Proceedings of the Royal Society of London B: Biological Sciences, 281(1797), 20141541.

Coelho, C. G., Falotico, T., Izar, P., Mannu, M., Resende, B. D. de, Siqueira, J., & Ottoni, E. B. (2015). Social learning strategies for nut-cracking by tufted capuchin monkeys (Sapajus spp.). Animal Cognition, 18(4), 911–919.

Coussi-Korbel, S., & Fragaszy, D. (1995). On the relation between social dynamics and social learning. Animal Behaviour, 50(6), 1441–1453. https://doi.org/10.1016/0003-3472(95)80001-8

de Vries, H. (1998). Finding a dominance order most consistent with a linear hierarchy: A new procedure and review. Animal Behaviour, 55, 827–843.

Drea, C. M., & Wallen, K. (1999). Low-status monkeys “play dumb” when learning in mixed social groups. Proceedings of the National Academy of Sciences, 96(22), 12965–12969. https://doi.org/10.1073/pnas.96.22.12965

Duboscq, J., Romano, V., MacIntosh, A., & Sueur, C. (2016). Social Information Transmission in Animals: Lessons from Studies of Diffusion. Frontiers in Psychology, 7. https://doi.org/10.3389/fpsyg.2016.01147

Duffy, G. A., Pike, T. W., & Laland, K. N. (2009). Size-dependent directed social learning in nine-spined sticklebacks. Animal Behaviour, 78(2), 371–375.

Farine, D. R. (2013). Animal social network inference and permutations for ecologists in R using asnipe. Methods in Ecology and Evolution, 4(12), 1187–1194. https://doi.org/10.1111/2041-210X.12121

Fragaszy, D., & Perry, S. E. (2003). Towards a Biology of Traditions. In D. Fragaszy & S. E. Perry (Eds.), The Biology of Traditions: Models and Evidence. Cambridge University Press.

Franz, M., & Nunn, C. L. (2009). Network-based diffusion analysis: A new method for detecting social learning. Proceedings of the Royal Society B: Biological Sciences. https://doi.org/10.1098/rspb.2008.1824

Galef, B. G., & Laland, K. N. (2005). Social Learning in Animals: Empirical Studies and Theoretical Models. BioScience, 55(6), 489–499. https://doi.org/10.1641/0006-3568(2005)055[0489:SLIAES]2.0.CO;2

Granovetter, M. S. (1977). The strength of weak ties. In Social networks (pp. 347–367). Elsevier.

Harcourt, J. L., Biau, S., Johnstone, R., & Manica, A. (2010). Boldness and information use in three-spined sticklebacks. Ethology, 116(5), 440–447.

Hobaiter, C., Poisot, T., Zuberbühler, K., Hoppitt, W., & Gruber, T. (2014). Social Network Analysis Shows Direct Evidence for Social Transmission of Tool Use in Wild Chimpanzees. PLOS Biology, 12(9), e1001960. https://doi.org/10.1371/journal.pbio.1001960

Hoppitt, W., Boogert, N. J., & Laland, K. N. (2010). Detecting social transmission in networks. Journal of Theoretical Biology, 263(4), 544–555.

Hoppitt, W., & Laland, K. N. (2013). Social learning: An introduction to mechanisms, methods, and models. Princeton University Press.

Horner, V., Proctor, D., Bonnie, K. E., Whiten, A., & de Waal, F. B. (2010). Prestige Affects Cultural Learning in Chimpanzees. PLOS ONE, 5(5), e10625. https://doi.org/10.1371/journal.pone.0010625

Huffman, M. A., & Hirata, S. (2003). Biological and ecological foundations of primate behavioral tradition. In D. Fragaszy & S. E. Perry (Eds.), The biology of traditions: Models and evidence (pp. 267–296).

Jones, T. B., Aplin, L. M., Devost, I., & Morand-Ferron, J. (2017). Individual and ecological determinants of social information transmission in the wild. Animal Behaviour, 129, 93–101.

Kameda, T., & Nakanishi, D. (2002). Cost–benefit analysis of social/cultural learning in a nonstationary uncertain environment: An evolutionary simulation and an experiment with human subjects. Evolution and Human Behavior, 23(5), 373–393. https://doi.org/10.1016/S1090-5138(02)00101-0

Katzir, G. (1983). Relationships between social structure and response to novelty in captive jackdaws, Corvus monedula L. II. Response to novel palatable food. Behaviour, 87(3–4), 183–208.

Kendal, R. L., Hopper, L. M., Whiten, A., Brosnan, S. F., Lambeth, S. P., Schapiro, S. J., & Hoppitt, W. (2015). Chimpanzees copy dominant and knowledgeable individuals: Implications for cultural diversity. Evolution and Human Behavior, 36(1), 65–72.

Krueger, K., Farmer, K., & Heinze, J. (2014). The effects of age, rank and neophobia on social learning in horses. Animal Cognition, 17(3), 645–655.

Kurvers, R. H., Prins, H. H., van Wieren, S. E., van Oers, K., Nolet, B. A., & Ydenberg, R. C. (2010). The effect of personality on social foraging: Shy barnacle geese scrounge more. Proceedings of the Royal Society B: Biological Sciences, 277(1681), 601–608.

Lerman, K., & Ghosh, R. (2010). Information contagion: An empirical study of the spread of news on Digg and Twitter social networks. Icwsm, 10, 90–97.

Luncz, L. V., & Boesch, C. (2014). Tradition over trend: Neighboring chimpanzee communities maintain differences in cultural behavior despite frequent immigration of adult females. American Journal of Primatology, 76(7), 649–657. https://doi.org/10.1002/ajp.22259

Marais, E. N. (1969). The soul of the ape. Atheneum.

Marshall, H. H., Carter, A. J., Coulson, T., Rowcliffe, J. M., & Cowlishaw, G. (2012). Exploring foraging decisions in a social primate using discrete-choice models. The American Naturalist, 180(4), 481–495.

McGrew, W. C. (1992). Chimpanzee material culture: Implications for human evolution. Cambridge University Press.

Mckerracher, L., Collard, M., & Henrich, J. (2015). The expression and adaptive significance of pregnancy-related nausea, vomiting, and aversions on Yasawa Island, Fiji. Evolution and Human Behavior, 36(2), 95–102.

Nishida, T. (1987). Local traditions and cultural transmission. In B. B. Smuts, D. L. Cheney, R. M. Seyfarth, R. W. Wrangham, & T. T. Struhsaker (Eds.), Primate societies. The University of Chicago Press.

Nishida, T., Matsusaka, T., & McGrew, W. C. (2009). Emergence, propagation or disappearance of novel behavioral patterns in the habituated chimpanzees of Mahale: A review. Primates, 50(1), 23–36. https://doi.org/10.1007/s10329-008-0109-y

Noldus Information Technology. (2003). Matman (1.1.4). Noldus Information Technology.

Nunn, C. L., Thrall, P. H., Bartz, K., Dasgupta, T., & Boesch, C. (2009). Do transmission mechanisms or social systems drive cultural dynamics in socially structured populations? Animal Behaviour, 77(6), 1515–1524.

Pereira, M. E., & Fairbanks, L. A. (2002). Juvenile primates: Life history, development and behavior, with a new foreword. University of Chicago Press.

Perry, S. E. (2009). Conformism in the food processing techniques of white-faced capuchin monkeys (Cebus capucinus). Animal Cognition, 12(5), 705–716. https://doi.org/10.1007/s10071-009-0230-3

Perry, S. E., Barrett, B. J., & Godoy, I. (2017). Older, sociable capuchins (Cebus capucinus) invent more social behaviors, but younger monkeys innovate more in other contexts. Proceedings of the National Academy of Sciences, 114(30), 7806–7813.

Perry, S. E., & Manson, J. H. (2003). Traditions in monkeys. Evolutionary Anthropology: Issues, News, and Reviews, 12(2), 71–81. https://doi.org/10.1002/evan.10105

Post, D. G., Hausfater, G., & McCuskey, SA. (1980). Feeding Behavior of Yellow Baboons (Papio cynocephalus): Relationship to Age, Gender and Dominance Rank. Folia Primatologica, 34(3–4), 170–195. https://doi.org/10.1159/000155954

Rapaport, L. G., & Brown, G. R. (2008). Social influences on foraging behavior in young nonhuman primates: Learning what, where, and how to eat. Evolutionary Anthropology: Issues, News, and Reviews, 17(4), 189–201.

Sapolsky, R. M., & Share, L. J. (2004). A Pacific Culture among Wild Baboons: Its Emergence and Transmission. PLOS Biology, 2(4), e106. https://doi.org/10.1371/journal.pbio.0020106

Schwab, C., Bugnyar, T., Schloegl, C., & Kotrschal, K. (2008). Enhanced social learning between siblings in common ravens, Corvus corax. Animal Behaviour, 75(2), 501–508.

Strum, S. C. (1975). Primate Predation: Interim Report on the Development of a Tradition in a Troop of Olive Baboons. Science, 187(4178), 755. https://doi.org/10.1126/science.187.4178.755

Strum, S. C. (1983). Baboon cues for eating meat. Journal of Human Evolution, 12(4), 327–336. https://doi.org/10.1016/S0047-2484(83)80159-6

Thornton, A., & Malapert, A. (2009). Experimental evidence for social transmission of food acquisition techniques in wild meerkats. Animal Behaviour, 78(2), 255–264.

van de Waal, E., Borgeaud, C., & Whiten, A. (2013). Potent Social Learning and Conformity Shape a Wild Primate’s Foraging Decisions. Science, 340(6131), 483. https://doi.org/10.1126/science.1232769

van de Waal, E., Renevey, N., Favre, C. M., & Bshary, R. (2010). Selective attention to philopatric models causes directed social learning in wild vervet monkeys. Proceedings of the Royal Society B: Biological Sciences. https://doi.org/10.1098/rspb.2009.2260

van Schaik, C. P. (2003). Local traditions in orangutans and chimpanzees: Social learning and social tolerance. In D. Fragaszy & S. E. Perry (Eds.), The Biology of Traditions: Models and Evidence. Cambridge University Press.

Voelkl, B., & Noë, R. (2010). Simulation of information propagation in real-life primate networks: Longevity, fecundity, fidelity. Behavioral Ecology and Sociobiology, 64(9), 1449–1459.

Watson, S. K., Vale, G. L., Hopper, L. M., Dean, L. G., Kendal, R. L., Price, E. E., Wood, L. A., Davis, S. J., Schapiro, S. J., Lambeth, S. P., & others. (2018). Chimpanzees demonstrate individual differences in social information use. Animal Cognition, 21(5), 639–650.

Weng, L., Menczer, F., & Ahn, Y.-Y. (2013). Virality prediction and community structure in social networks. Scientific Reports, 3, 2522.

Whitehead, H. (2008). Analyzing animal societies: Quantitative methods for vertebrate social analysis. University of Chicago Press.

Whiten, A., & Mesoudi, A. (2008). Establishing an experimental science of culture: Animal social diffusion experiments. Philosophical Transactions of the Royal Society B: Biological Sciences, 363(1509), 3477. https://doi.org/10.1098/rstb.2008.0134

Wood, L. A., Kendal, R. L., & Flynn, E. G. (2013). Whom do children copy? Model-based biases in social learning. Developmental Review, 33(4), 341–356.

